# Mucosal and systemic responses to SARS-CoV-2 vaccination in infection naïve and experienced individuals

**DOI:** 10.1101/2021.12.13.472159

**Authors:** Mohammad M. Sajadi, Amber Myers, James Logue, Saman Saadat, Narjes Shokatpour, James Quinn, Michelle Newman, Megan Deming, Zahra Rikhtegaran Tehrani, Maryam Karimi, Abdolrahim Abbasi, Mike Shlyak, Matthew B. Frieman, Shane Crotty, Anthony D. Harris

**Author notes:** Corresponding author: Mohammad Sajadi, MD Institute of Human Virology, Global Virus Network Center of Excellence, University of Maryland School of Medicine, 725 W. Lombard St. (N548), Baltimore, MD 21201.

## Abstract

With much of the world infected with or vaccinated against SARS-CoV-2, understanding the immune responses to the SARS-CoV-2 spike (S) protein in different situations is crucial to controlling the pandemic. We studied the clinical, systemic, mucosal, and cellular responses to two doses of SARS-CoV-2 mRNA vaccines in 62 individuals with and without prior SARS-CoV-2 exposure that were divided into three groups based on serostatus and/or degree of symptoms: Antibody negative, Asymptomatic, and Symptomatic. In the previously SARS-CoV-2-infected (SARS2-infected) Asymptomatic and Symptomatic groups, symptoms related to a recall response were elicited after the first vaccination. Anti-S trimer IgA and IgG levels peaked after 1^st^ vaccination in the SARS2-infected groups, and were higher that the in the SARS2-naive group in the plasma and nasal samples at all time points. Neutralizing antibodies titers were also higher against the WA-1 and B.1.617.2 (Delta) variants of SARS-CoV-2 in the SARS2-infected compared to SARS2-naïve vaccinees. After the first vaccination, differences in cellular immunity were not evident between groups, but the AIM^+^ CD4^+^ cell response correlated with durability of humoral immunity against the SARS-CoV-2 S protein. In those SARS2-infected, the number of vaccinations needed for protection, the durability, and need for boosters are unknown. However, the lingering differences between the SARS2-infected and SARS2-naïve up to 10 months post-vaccination could explain the decreased reinfection rates in the SARS2-infected vaccinees recently reported and suggests that additional strategies (such as boosting of the SARS2-naïve vaccinees) are needed to narrow the differences observed between these groups.

## INTRODUCTION

Although several recent studies have shown protection from reinfection amongst those with SARS-CoV-2 infection who are subsequently vaccinated (*1, 2*), controversy exists as to the degree of protection from reinfection in those with prior COVID-19 infection, as well as the role for vaccination and boosters in this population. Understanding the primary immune responses to natural infection and vaccination, as well as the secondary responses in those previously infected or vaccinated is key to understanding the protection. Given the recent findings of a decrease in durability of protection after vaccination (*3*), this has become even more urgent. The current study was undertaken to investigate the primary responses in COVID-19 naïve vaccinees, as well as secondary responses in those with previous COVID-19. To our knowledge, no group has comprehensively studied these responses to include clinical symptoms, circulating binding and neutralization, and cellular and mucosal immunity.

## RESULTS

### First vaccination elicits elevated symptoms in SARS2-infected individuals

A total of 3816 health care workers (HCW) were enrolled in the serosurvey study (*4*), of which 151 were randomly contacted, and 67 volunteers enrolled. Of the 67, 62 had received both vaccine doses (4 had dropped out prior to second vaccine and one was diagnosed with COVID-19 before second vaccine). The 62 who received both vaccines divided into the following groups: 19 Antibody negative, 17 Asymptomatic antibody positive, and 26 Symptomatic antibody positive (Table 1).

**Table 1.** Study Population Baseline Characteristics.

|  |  | **Ab negative**<br>**(n=19)** | **Asymptomatic**<br>**(n=17)** | **Symptomatic**<br>**(n=26)** |
| --- | --- | --- | --- | --- |
| Age (years) | Median | 42 | 39 | 38 |
|  | Range | 29-62 | 25-72 | 23-59 |
| Sex [n (%)] | Male | 5 (26) | 4 (24) | 3 (12) |
|  | Female | 14 (74) | 13 (76) | 23 (88) |
| Race/Ethnicity [n (%)]* | Black or African American | 2 (11) | 5 (29) | 5 (19) |
|  | White or Caucasian | 14 (74) | 9 (53) | 18 (69) |
|  | Asian | 3 (16) | 3 (18) | 3 (14) |
|  | Hispanic or Latino | 0 (0) | 0 (0) | 0 (0) |
| Vaccine [n (%)] | Pfizer | 11 (58) | 7 (41) | 13 (50) |
|  | Moderna | 8 (42) | 10 (59) | 13 (50) |
| Days symptomatic | Median | N/A | 0 | 10 |
|  | Range | N/A | 0-2 | 3-34 |
| Hospitalization [n (%)] |  | N/A | 0 (0) | 4 (15) |
| Months from COVID PCR+ to vaccination | Median (percent PCR tested) | N/A | 9.0 (41%) | 8.0 (73%) |
|  | Range | N/A | 6.5-9.7 | 6.0-10.3 |
| Months from COVID IgG+ to vaccination | Median | N/A | 6.1 | 6.1 |
|  | Range | N/A | 4.8-7.1 | 3.8-7.2 |

N/A-not applicable \*-Race/ethnicity data derived from self-report. Of research interest because different race/ethnicity may react differently to vaccine.

After the first vaccination, individuals in the Asymptomatic and Symptomatic groups had higher rates of systemic symptoms compared to those in the Antibody negative group, for fever, chills, fatigue, myalgia, headache in the Symptomatic group (**Fig. 1**). After the second vaccination, the Ab negative had an increase in symptom reporting compared to the first vaccination, and any differences with the Symptomatic group were no longer statistically significant. However, after the second vaccination, the Asymptomatic Group had less fatigue, myalgias, and headaches compared to the Ab negative group (**Fig. 1**). After the first and second vaccination, the Asymptomatic group had less symptom reporting in nearly every category, and although the individual categories did not reach statistical significance, the Asymptomatic group had less systemic symptoms than the Symptomatic when considering the number of systemic symptoms reported in each category of 1 more (71% vs 96%), or 2 or more (41% vs 77%) systemic symptoms (P=.03 and P=.03, respectively).

**Figure 1.**
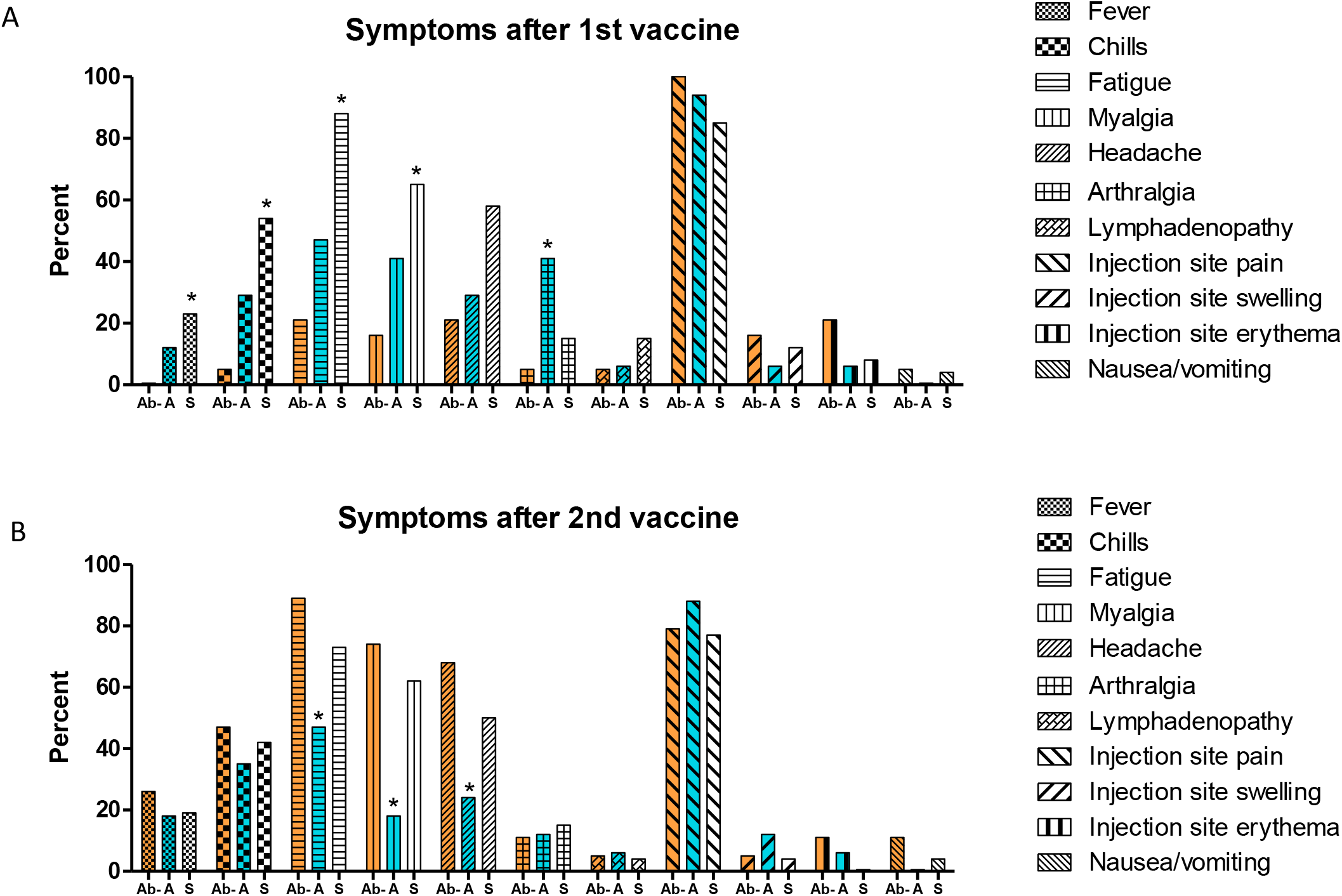
Reported symptoms after first and second vaccination. Percent of reported symptoms after first (A) and second (B) vaccinations shown on the Y axis. Ab negative, Asymptomatic, and Symptomatic groups shown in orange, blue, and white columns, respectively. Each symptom was assessed by 2-tailed Fisher’s exact test between the Ab negative and the Asymptomatic or Symptomatic groups, with P values <.05 being considered significant and shown with asterisks above the columns.

### SARS2-infected individuals have higher plasma and nasal antibodies compared to SARS2-naïve individuals

At baseline, the Antibody negative groups had undetectable endpoint titers of IgM, IgA, and IgG antibodies to the SARS-CoV-2 spike trimer, whereas these could be detected to varying degrees in those in the Asymptomatic and Symptomatic groups (**Fig. 2A**). IgM, IgA, and IgG S trimer endpoint titers continued to increase after 2^nd^ vaccination in the Ab negative group, while they plateaued after the first vaccination in the Asymptomatic and Symptomatic Groups (**Fig. 2A**). IgG S trimer endpoint titers after each vaccination were higher in the Asymptomatic and Symptomatic Groups compared to the Ab negative group (**Fig. 2A**), and this difference continued until the study completion, up to 10 months after the second vaccination (**Fig. 2B**). At the last time point the median endpoint titers for the Antibody negative group was 2,700, and statistically lower compared to the Asymptomatic (8,100) and Symptomatic (8,100) groups (P =.0005 and <.0001, respectively). There was no significant difference in the slope of the decline of IgG binding titers between 3 month after the second dose and up to 10 months after the second dose in the Ab negative (−0.12501 log titer/month) compared to the Asymptomatic (−0.09471 log titer/month) and Symptomatic (−0.08233 log titer/month) groups (P=0.37 and P=0.67, respectively) (**Fig. 2B**). Comparing Moderna versus Pfizer recipients within each group, we noted larger IgG anti-S binding responses in SARS2-infected volunteers14 days after the first Moderna vaccine (P= .0056), but not at any other time points (fig S1A).

**Figure 2.**
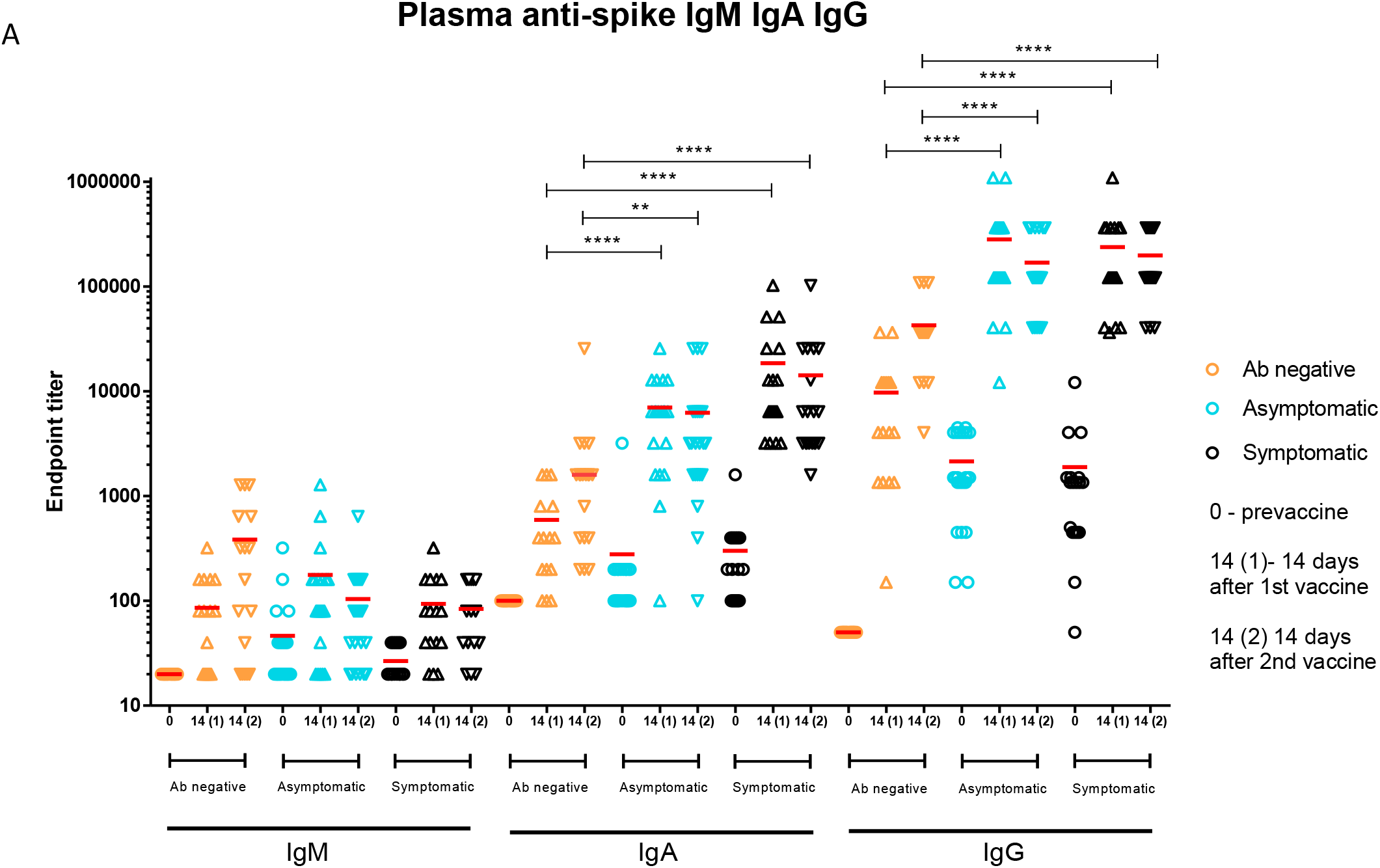
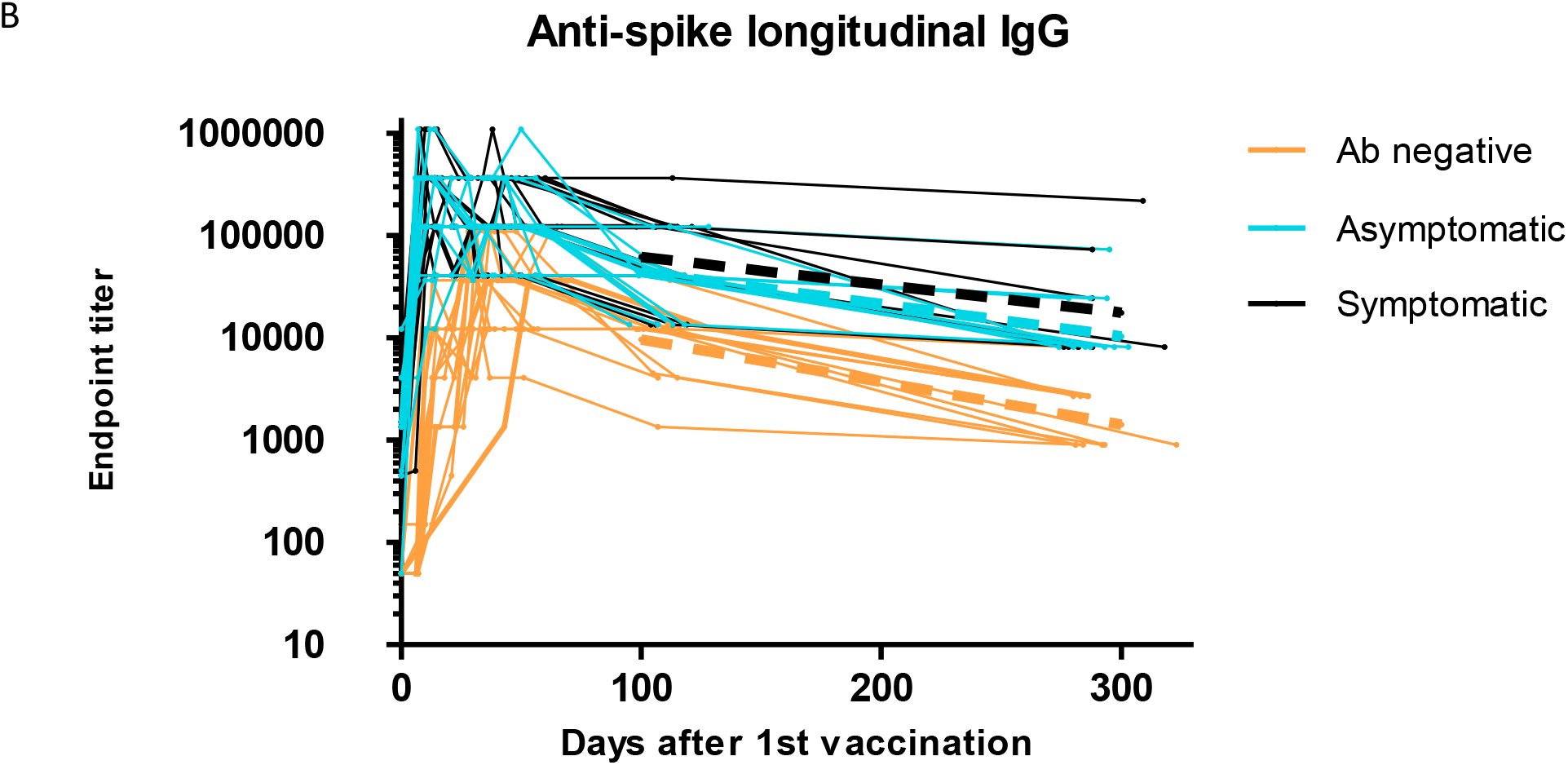
IgM, IgA, and IgG ELISA responses to SARS-CoV-2 S trimer. A) IgM, IgA, and IgG endpoint titers were measured at baseline, and at 14 days after 1^st^ and second 2^nd^ vaccination. Horizontal red lines represent median values. B). IgG endpoint titers measured at all time points until 10 months after second vaccination. Dotted lines represent decay slopes for each cohort between months 3 and 10. Ab negative, Asymptomatic, and Symptomatic groups shown in orange, blue, and black colors, respectively. Y-axis represents reciprocal endpoint titer to SARS-CoV-2 S trimer in a logarithmic scale. * = P ≤ 0.05; ** = P ≤ 0.01; *** = P ≤ 0.001; **** = P ≤ 0.0001

The nasal responses to the S trimer largely mirrored the serum responses. The Ab negative group showed sequential increases in nasal IgA and IgG after the two vaccines. The Asymptomatic and Symptomatic groups show peaks at the 14 day time point after the first vaccine with no further increased after second vaccination (**Fig. 3**). In the SARS2-infected groups, the IgG levels went up 81-90 fold compared to baseline, while the IgA levels went up 32-64 fold compared to baseline.

**Figure 3.**
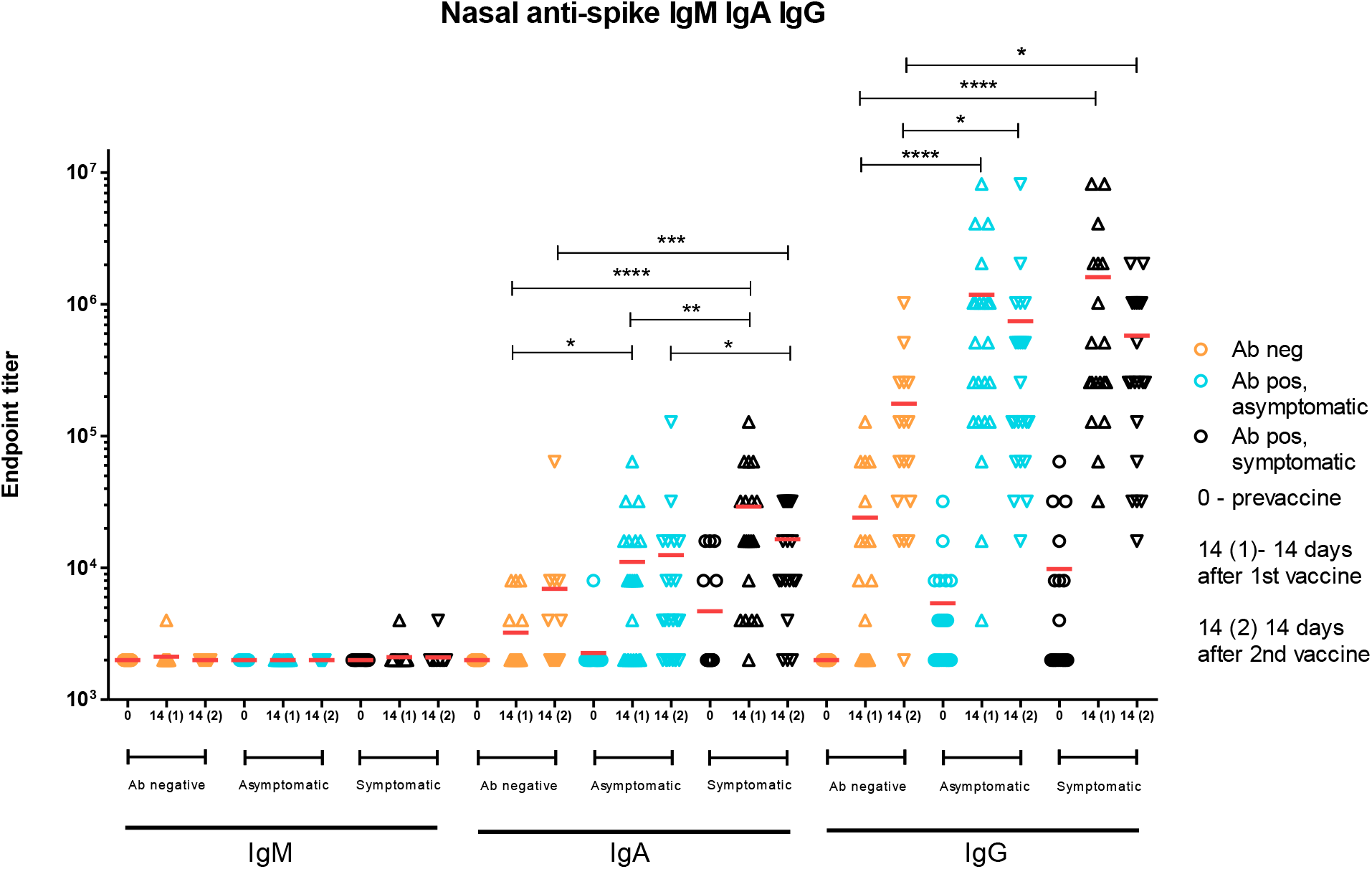
Nasal sample ELISAs and IgG/IgA ratios in nasal and plasma samples. IgA and IgG endpoint titers in nasal samples were measured at baseline, and at 14 days after 1^st^ and second 2^nd^ vaccination. Horizontal red lines represent median values. * = P ≤ 0.05; ** = P ≤ 0.01; *** = P ≤ 0.001; **** = P ≤ 0.0001

In the nasal samples the median IgG/IgA ratio for total non-vaccine specific antibody was 0.76 compared to 7.97 in the plasma (P<.0001) and at baseline in the Asymptomatic and Symptomatic groups the median SARS-CoV-2 specific IgG/IgA ratio was 2.0 compared to 7.5 in the plasma (P<.0001) (fig. S2A). After the first vaccination the median SARS-CoV-2 specific IgG/IgA increased to 64 in the nasal samples compared to 23.73 in the plasma (P=.04) (fig. S2A). Spike specific antibody carrying the secretory component was not seen at baseline but was detected in a majority of those in the Asymptomatic and Symptomatic groups after the first vaccination (fig S2B).

The plasma samples were tested against the WA-1 and B.1.617.2 (Delta) variants of SARS-CoV-2. In the Ab negative group the neutralization titers for both Wuhan and Delta variants sequentially increased until 14 days after the second vaccination (**Fig. 4A and B**). In the Asymptomatic and Symptomatic groups, the endpoint titers peaked at 14 days after the first vaccination, and remained stable before and after the second vaccination. By 14 days after the second vaccination, median ID99 WA.1 neutralization titers were 5,120 in the Ab negative group, compared to the 51,200 in the Asymptomatic and 81,920 in the Symptomatic groups (P<.0001 for each) (**Fig. 4A**). Comparing Moderna versus Pfizer recipients within each group, we noted larger neutralization responses in SARS2-infected volunteers against WA-1 14 days after the first vaccine (P=.004), but not at any other time points (fig. S1E).

**Figure 4.**
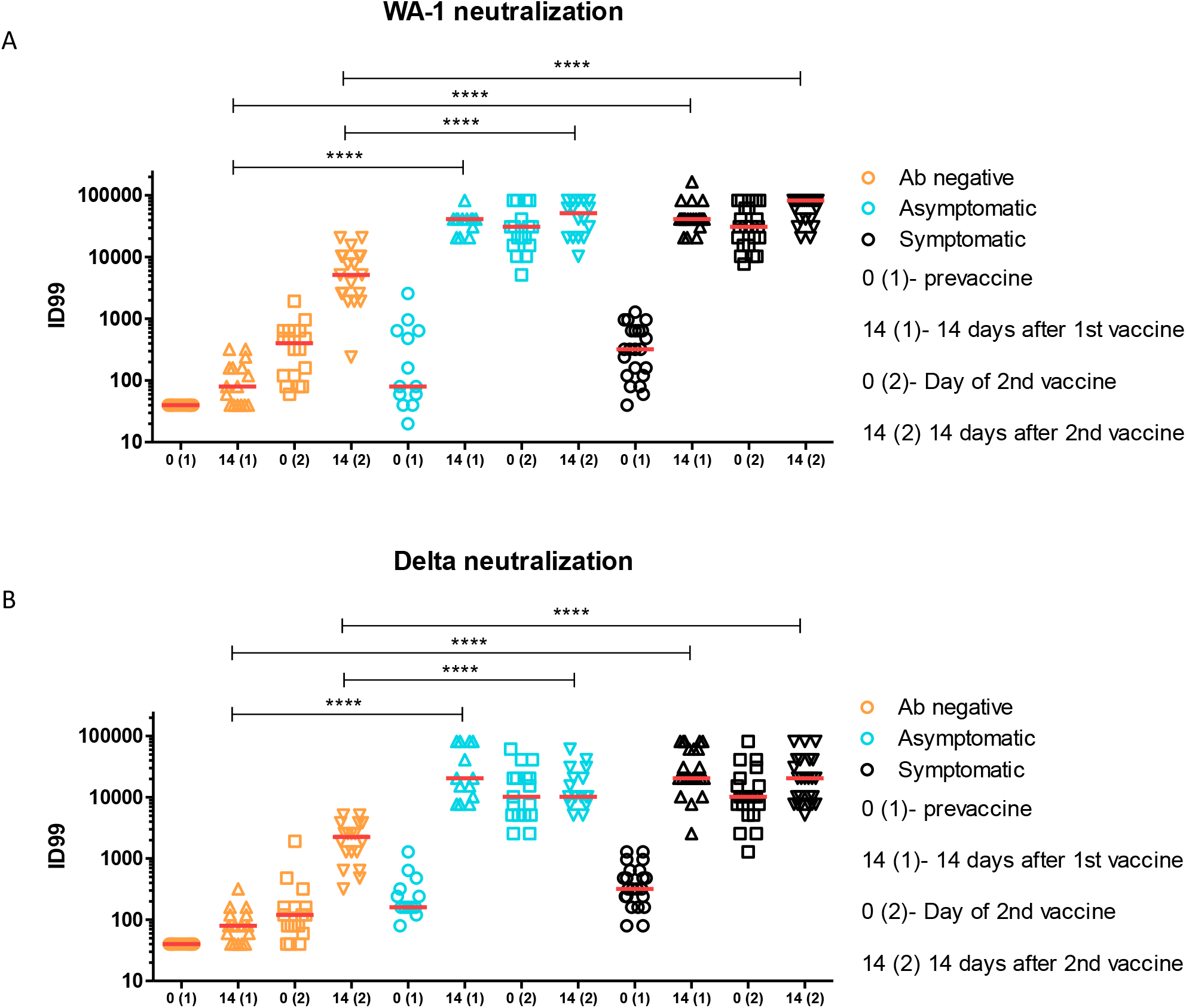
Live virus plasma neutralization. Live virus neutralization in plasma against WA-1 (A) and B.1.617.2 (“Delta”) (B) measured at baseline, and at 14 days after 1^st^ vaccination, and prior to and 14 days after the second 2^nd^ vaccination. ID99 defined as highest dilution at which 99% cells were protected. Horizontal red lines represent median values. * = P ≤ 0.05; ** = P ≤ 0.01; *** = P ≤ 0.001; **** = P ≤ 0.0001

### After 1^st^ vaccination cellular responses similar in all groups

We compared the CD4^+^ and CD8^+^ T cell reactivity of the three cohorts against SARS-CoV-2 S by activation induced cell marker (AIM) and intracellular cytokine staining (ICS) assays (**Fig. 5A, C, E, H**, and figs. S3 and S4). When comparing the AIM responses at baseline, the Symptomatic group had significantly higher levels of CD4 and cT_FH_ responses to Spike compared to the Antibody negative group (P<.0001 and P=.009, respectively) (**Fig. 5B and D**). The antibody negative group saw increases in CD4 and cT_FH_ AIM responses after each vaccination, while AIM responses in the Asymptomatic and Symptomatic groups peaked after the first vaccination (**Fig. 5B and D**). There was no statistically significant difference between the AIM responses to CD4 or cT_FH_ between the SARS2-naive and infected groups after the 1^st^ or 2^nd^ vaccinations (**Fig. 5B and D**).

**Figure 5.**
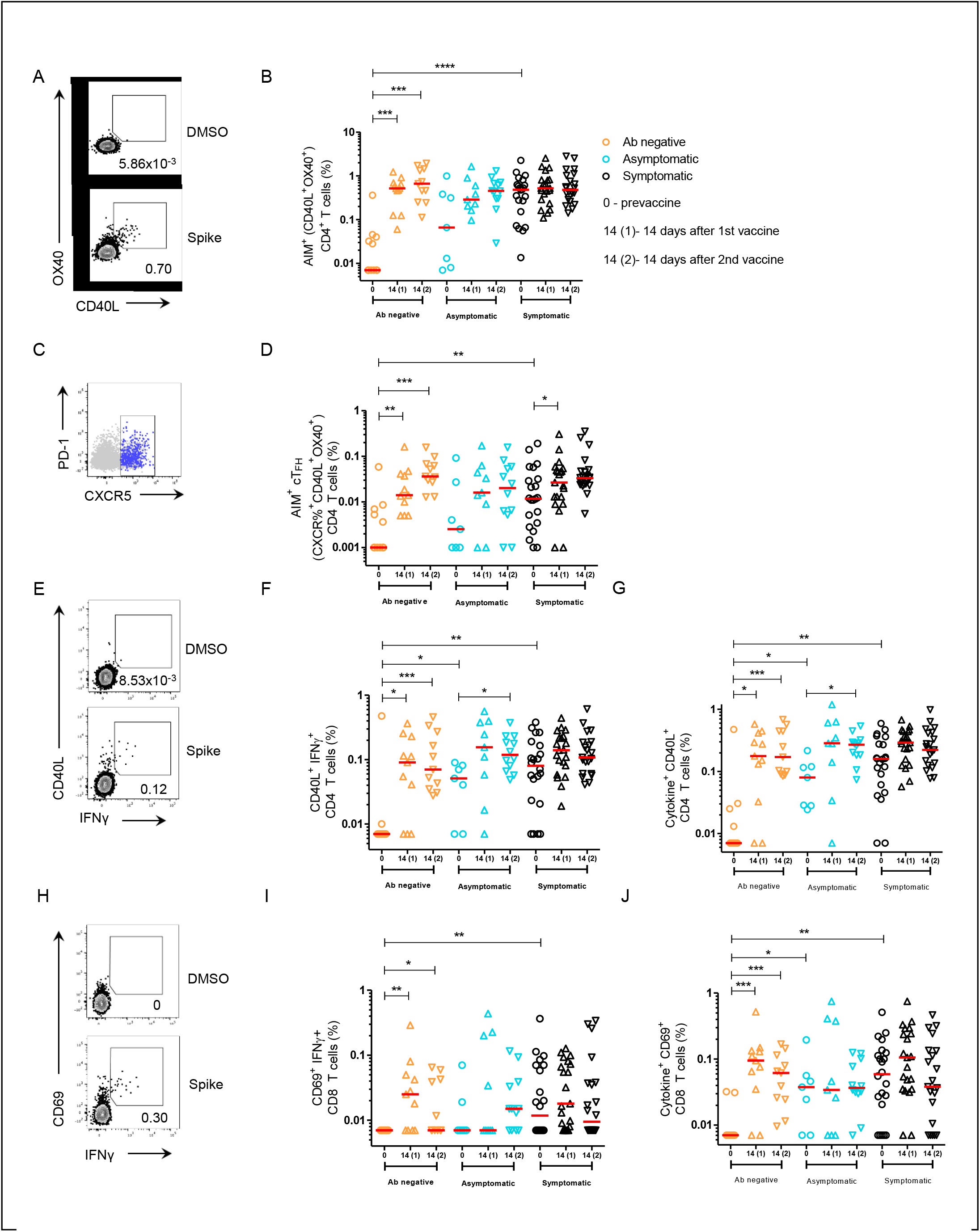
SARS-CoV-2 S-specific CD4+ and CD8+ T cells following vaccination in unexposed and previously infected individuals. (A) Representative gating strategy of AIM+ (surface CD40L+ OX40+) CD4+ T cells stimulated with S peptide pool or unstimulated (DMSO). (B) AIM+ S-specific CD4+ T cells of total CD4+ T cells. (C) Representative gating strategy of AIM+ (CD40L+ OX40+) circulating T follicular helper (cT_FH_) cells (Blue, CXCR5+) overlaid on total CD4+ T cells (Gray). (D) AIM+ S-specific cT_FH_ cells of total CD4+ T cells. (E) Representative gating strategy of CD40L+ IFNγ+ CD4+ T cells. (F) Spike-specific CD40L+ IFNγ + CD4+ T cells of total CD4+ T cells. (G) Spike-specific CD40L+ CD4+ T cells producing IFNγ, IL-2 or TNFα of total CD4+ T cells. (H) Representative gating strategy of CD69+ IFNγ+ CD8+ T cells. (I) Spike-specific CD69+ IFNγ + CD8+ T cells of total CD8+ T cells. (J) Spike-specific CD69+ CD8+ T cells producing IFNγ, IL-2 or TNFα of total CD8+ T cells. All data are background subtracted. Error bars represent geometric mean and geometric standard deviation. Horizontal red lines represent median values. * = P ≤ 0.05; ** = P ≤ 0.01; *** = P ≤ 0.001;

In the CD4^+^ ICS assay, significantly lower S-specific IFNγ and cytokine staining were seen in the Antibody negative group at baseline compared to the Asymptomatic (P=.04 and P =.01, respectively) and Symptomatic (P=.003 and P=.001, respectively) groups (**Fig. 5F-G)**. Compared to baseline levels, after 1^st^ vaccination peak percentages of IFNγ and cytokine secreting cells significantly increase in the Antibody negative group only (P=.02 and P=.02, respectively). Second vaccination did not see further increases in any of the groups (**Fig. 5F-G)**.

S-specific CD8^+^ T cell IFNγ and cytokine expression differences were noted between the Symptomatic and Antibody negative groups at baseline (P=.006 and P=.002, respectively) (**Fig. 5I and J)**. Compared to baseline levels, after 1^st^ vaccination peak percentages of IFNγ and cytokine secreting cells significantly increase in the Antibody negative group only (P=.003 and P=.001, respectively). (**Fig. 5I and J**). The percentage of IFNγ and cytokine secreting cells in the Asymptomatic and Symptomatic groups did not significantly change before or after vaccinations (**Fig. 5I and J**). There was a strong positive correlation between the AIM^+^ CD4^+^ cells and percentage of IFNγ secreting CD4^+^ cells in each of the Antibody negative (P< .0001), Asymptomatic (P=.003), and Symptomatic (P< .0001) groups (fig S5). In the Antibody negative group, Moderna vaccines recipients had statistically significant higher S-specific CD4^+^ T cell and cT_FH_ cells detected by AIM assay, and higher S-specific CD4^+^ T cell and CD8^+^ T cell ICS responses than Pfizer recipients 14 days after the second vaccine (figs S6 and S7).

The cellular and antibody data were analyzed for correlations. In the SARS2-infected groups, there were positive correlations between S-specific CD4^+^ cells (Aim^+^) with S IgG titers pre-vaccination (P=.01), at month 3 (P=.03) and Month 10 (P=.006) (fig. S8A, I, K). In the SARS2-infected groups, there were also positive correlations between AIM^+^ cT_FH_ cells with S IgG titers at Month 10 (P=.04) (fig. S8L). There were positive correlations between AIM^+^ CD4^+^ cells with neutralization titers to WA.1 pre-vaccination (P=.047), and Day 14 after second vaccination (P=.003) (fig. S9A, E).

## DISCUSSION

Several years into the COVID-19 pandemic, key questions about the immune response to SARS-CoV-2 S protein after natural infection and vaccination remain. Currently, waning mRNA vaccine immunity has been demonstrated in a number of studies (*3, 5-8*), and thus understanding the primary and secondary responses to SARS-CoV-2 is imperative. In this study, we had the opportunity to evaluate primary responses in SARS2-naïve vaccinees, as well as secondary responses in the SARS2-infected vaccinees. The primary and secondary responses had a clear set of differences including symptoms and antibody levels.

After the first dose of vaccine, those with a history of SARS2 infection had more symptoms, consistent with a secondary immune response. Interestingly, those in the Symptomatic group also had a higher percentage of symptoms compared to the Asymptomatic group. Given the lack of significant immune differences observed between the two groups, this opens the possibility that the elevated secondary response to vaccination could be related to exaggerated response to antigens in this group (genetic differences), though it can also be a reflection of a higher viral burden during SARS2-infection (consequence of having more severe COVID-19).

Prior to vaccination, the SARS2-infected groups had elevated levels of AIM^+^ CD4 and CD8 cells directed at the S protein of SARS-CoV-2, cells that contributed to the secondary response after vaccination. Within the SARS2-infected groups, those in the Symptomatic group tended to have slightly higher baseline levels than the Asymptomatic group. By 14 days after the first vaccination, the cellular responses in all 3 groups reached a peak, with no apparent changes at day 14 after the second vaccination.

Two weeks after the vaccination series, IgM responses were higher in the Ab negative group, who were mounting a primary response, compared to the SARS2-infected groups. However, IgA and IgG responses were higher at all time points measured, and up to 10 months in the case of IgG, in the SARS2-infected groups. The higher antibody levels in the SARS2-infected was also true of neutralizing antibody against the WA-1 and Delta variants of SARS-CoV-2, which were approximately 10 times higher in the SARS2-infected groups two weeks after the second vaccination.

The findings of higher binding and neutralization titers in vaccinees with prior COVID-19 indicative of a secondary immune response has been noted in other studies as well (*9-19*). We have previously shown that peaking titers in the Symptomatic and Asymptomatic groups occur by Day 7 after the first vaccination (*13*), while peak titers in the Ab negative group occurs after the second vaccination. Two recent studies have documented protection from infection in vaccinees with prior COVID-19 (*1, 2*); however, it remains unknown if 2 doses rather than 1 dose of an mRNA vaccine is needed for protection. Importantly, one of these studies has shown that SARS2-infected vaccinees appear to have less reinfection than SARS2-naïve vaccinees (*2*), and the finding of elevated binding and neutralization titers in the former group could explain the increased protection.

This study was continued 10 months after the second vaccination, so there was an opportunity to study the early durability of the responses in the different groups. There was no difference in the slope of decline between the groups, but the IgG antibody levels in the SARS2-infected groups remained higher at each time point, including at last time point. Interestingly, the AIM+ CD4^+^ cell response at baseline in the SARS2-infected correlated with IgG anti-S antibody responses, but only at baseline, 3 months, and 9-10 months. Given the lack of correlation with peak titers, this suggests that AIM^+^ CD4^+^ response tracks with the durability rather than the magnitude of the IgG response. Neutralizing antibody also correlated with AIM^+^ CD4^+^ cells at baseline and 14 days after vaccinations, but later time points were not assessed for neutralization, and so it is unknown if neutralizing antibody correlations with AIM^+^ CD4^+^ cells would differ in the late time points. Currently, there is a lack of data regarding the level of protective antibodies and late vaccine efficacy in those with prior SARS2-infection, but the data generated in this study can help inform those when available. Other questions that remain unanswered are the role of booster vaccination in the SARS2-infected population, and if boosting the SARS2-naïve vaccine recipients will equalize the antibody response and peak titers amongst all the groups.

Mucosal immunity is an understudied but important facet of COVID-19 humoral immunity, as the respiratory mucosa are the first site of contact between SARS-CoV-2 and humans. In the nasal mucosa, it is known that IgA as well as IgG are abundant with IgA predominating (*20*), results that were seen in the non-vaccine specific nasal responses in this study. Most of the IgA is thought to be secretory, in the form of dimers, and produced locally, whereas IgG produced in the bone marrow or lymph nodes arrives by a process of transcytosis involving FcRn receptor (*21*). Anti-S trimer IgA and IgG were not detected in all SARS2-infected volunteers at baseline, but this may be due to the fact that swabs were placed in 1ml of PBS, and thus diluted 1000 times what their concentration would be on the mucosal surface prior to testing. Nevertheless, S-specific IgG/IgA ratio in the SARS2-infected individuals pre-vaccination was much lower in the nose than blood, suggesting local IgA production. After vaccination, anti-S IgG titers in the nasal mucosa outstrip the IgA, as reflected in the skewed IgG/IgA ratios, some of which exceeded 100:1. These changes mirrored the concentrations measured in plasma, whereby IgG levels outpaced IgA. While we did measure antibody attached to secretory protein in the nasal samples (presumably IgA because IgM was undetectable in all but a handful of samples), further studies needs to be carried to definitively identify the type and source of these antibodies. Nevertheless, the presence of both IgG and IgA at the site of first viral contact is important in that they are likely involved in the process by which the vaccines afford protection from acquisition and subsequent disease. Although plasma IgG binding and neutralizing antibody has been the focus of correlative studies (*22, 23*), this is currently not definitely proven and IgA may play an important role as well. Additionally, the importance of each could differ based on whether an individual was infected or vaccinated, as local IgA production after infection may be a significant contributor to mucosal immunity compared to vaccination where most of the antibodies arrive from the systemic circulation.

The two vaccines used in this study, Moderna mRNA-1273 and Pfizer BNT162b2, share similarities in design and mode of action; however, they differ in the dosage administered, with the Moderna using 3.3 times as much (100ug vs 30ug). We noted differences in cellular responses in the SARS2-naïve group 2 weeks after the second vaccination in all CD4, cT_FH_, and CD8 assays that we analyzed, with the Moderna group having elevated responses. By contrast, in the SARS2-infected group, only cT_FH_ responses were noted to be elevated in the Moderna recipients in the Symptomatic group 2 weeks after first vaccination.

The strengths of this study include using a clearly defined cohort with regimented sampling points. Additional strengths include use of vaccinees who received Moderna or Pfizer, as well as studying clinical, cellular, and mucosal parameters. This study was limited by sample size, and thus a larger sample size could have found additional differences between the groups based on prior SARS-CoV-2 exposure, type of vaccination, or other factors. Finally, although the studies undertaken were fairly comprehensive, some key sites involved in immune regulation (lymph nodes, lungs) were not involved in the current study.

This study found key differences in the response to SARS-CoV-2 vaccination depending on previous exposure to the virus. SARS2-infected individuals had higher binding and neutralization titers throughout the study. This was also true of the relevant classes of anti-S antibody (IgG and IgA) at mucosal sites which are likely involved in the protective efficacy of the mRNA vaccines. Finally, while differences in cellular immunity were less evident between groups after vaccination, there was data supporting that the AIM+ CD4^+^ cell response could be tied to durability of humoral immunity against the SARS-CoV-2 S protein. In those with prior SARS2 infection, the number of vaccinations needed for protection, the durability, and need for boosters are unknown; however, the lingering differences between the SARS2-infected and SARS2-naïve up to 10 months post-vaccination suggests that additional strategies (such as boosting of the SARS2-naïve vaccinees) are needed to narrow the immunological differences observed between the groups in this study, which are likely related to the

## MATERIALS AND METHODS

### Study Design

HCW who had previously enrolled in a hospital-wide serosurvey study in the summer of 2020 (*4*), conducted at the University of Maryland Medical Center, were randomly contacted based on stratification into three groups, as previously described (*13*): SARS-CoV-2 IgG antibody negative (Ab negative); IgG positive asymptomatic and minimally symptomatic COVID-19 (Asymptomatic); and IgG positive with history of symptomatic (≥ 3 days of symptoms) COVID-19 (Symptomatic). Participants were vaccinated with either the Pfizer-BioNTech or Moderna vaccine, depending on personal preference and availability. Blood was drawn at Day 0 (or baseline), 7, 10, and 14 after first vaccination, and 0, 7, 10, 14, and 28 days, as well as 3 and 10 months after second vaccination. Upon study enrollment, volunteers were given a questionnaire regarding COVID-19 history, and after vaccination they were given a questionnaire regarding post vaccination symptoms. This included presence of subjective fever, chills, fatigue, myalgia, headache, arthralgia, axillary lymphadenopathy, injection site pain, injection site swelling, injection site erythema, and nausea/vomiting.

### ELISA

IgA, IgM, IgG, and secretory component (SC) ELISA assays to SARS-CoV-2 S trimer were made by modifying an assay (*13, 24*) to give a read-out of end-point binding titers. IgM, IgA, and IgG responses were measured in the plasma, and nasal samples. Additionally, antibody responses to the secretory component (SC), indicative of secretory IgA or IgM, were evaluated in nasal and plasma samples. For nasal samples, the anterior nares were swabbed with a flock swab that was wetted with PBS prior to swabbing, and then placed in a tube with 1 ml of PBS.

### Neutralization Assay

Live virus neutralization assays were carried out, as previously described (*13, 25*), on plasma from Day 0 and 14 after the first vaccination, and Day 14 after second vaccination. Briefly, Serial dilutions of plasma was incubated for one hour with 200TCID50 of the WA-1 or B.1.617.2 variants of SARS-CoV-2. This admixture was added to Vero cells, and cytopathic effect was assayed visually. ID99 were recorded as the first plasma dilution with cytopathic effect similar to the virus-only control wells.

### Activation induced cell marker (AIM) assay

To quantify the S-specific T cell response, a S megapool (MP) containing overlapping peptides of the entire S protein sequence was used. In addition, MP remainder (MP_R) containing non-S epitopes in SARS-CoV-2 was used, as previously described (*26*). Peripheral blood mononuclear cells (PBMCs) were incubated with 1 μg/mL of S MP or MP_R for 24 hours in a 96-well U bottom plate at 1×106 cells/well in RPMI media containing 5% human serum AB (Gemini Bio), 1% GlutaMAX (Thermo Fisher Scientific) and 1% Penicillin-Streptomycin (Thermo Fisher Scientific). DMSO (0.1%) and staphylococcal enterotoxin B (SEB, 1 μg/mL) were used as negative and positive controls, respectively. Fifteen minutes prior to the addition of the MPs, 0.5 μg/mL anti-CD40 mAb (Miltenyi Biotec) and CXCR5 antibody were added. Following the 24 hour incubation, PBMCs were surface stained with antibodies in Table S1 for 30 mins at 4°C, washed in PBS, fixed for 10 mins in 4% formaldehyde (BD Biosciences Cytofix), washed in PBS and acquired on a BD FACSCelesta Cell Analyzer. AIM+ antigen-specific CD4+ T cells were defined as (surface CD40L+ OX40+) and antigen-specific circulating T follicular helper (cT_FH_) cells were defined as (CXCR5+ surface CD40L+ OX40+).

### Intracellular cytokine staining (ICS) assay

PBMCs were incubated with 1 μg/mL of S MP or MP_R for 24 hours in a 96-well U bottom plate at 1×10^6^ cells/well in RPMI media containing 5% human serum AB (Gemini Bio), 1% GlutaMAX (Thermo Fisher Scientific) and 1% Penicillin-Streptomycin (Thermo Fisher Scientific), as previously described (*27*). DMSO (0.1%) and staphylococcal enterotoxin B (SEB, 1 μg/mL) were used as negative and positive controls, respectively. Fifteen minutes prior to the addition of the MPs, 0.5 μg/mL anti-CD40 mAb (Miltenyi Biotec) and CXCR5 antibody were added. Following the 24 hour incubation, monensin (BD Biosciences GolgiStop) and brefeldin A (BD Biosciences GolgiPlug) were added and PBMCs were incubated for an additional 4 hours. PBMCs were surface stained with antibodies in Table S2 for 30 mins at 4°C, washed in PBS, and then fixed and permeabilized (1x BD Biosciences Cytofix/Cytoperm) for 30 mins. After fixation, PBMCs were washed in 1x Perm/Wash buffer (BD Biosciences) and stained for 45 mins at 4°C with the intracellular antibody panel in Table S3 and acquired on a BD FACSCelesta Cell Analyzer. Antigen-specific CD4+ T cells were defined as CD40L+ and IFNγ+, TNFα+, or IL-2+ and antigen-specific CD8+ T cells were defined as CD69+ and IFNγ+, TNFα+, or IL-2+.

### Statistical Analysis

The reciprocal end-point binding titers represent the maximal dilution of plasma that achieves binding. For T cell assays, sample quality was assessed by CD4+ AIM+ (CD40L+ OX40+) SEB response. Samples with a SEB CD4+ AIM+ response lower than one-half the median (<3.7% CD4+ AIM+ of total CD4+ T cells) were eliminated from the analysis. T cell data was analyzed using FlowJo 10.7.1. All data was background subtracted. Statistical analysis was carried out with GraphPad Prism 5 (GraphPad Software). Differences between groups were tested by the 2-tailed Fisher’s exact test or Mann-Whitney test, with a p<.05 being considered significant. Correlation graphs were analyzed using Spearman’s Correlation test. All volunteers in this study provided informed consent for the IRB approved study (University of Maryland, Baltimore).

## Acknowledgements

ADH supported by CDC grant U01CK000556-02-01. This work was supported in part by Merit Award # I01 BX005469-01 from the United States (U.S.) Department of Veterans Affairs Biomedical Laboratory Research and Development Service. The contents do not represent the views of the U.S. Department of Veterans Affairs or the United States Government.

---

**Supplementary Figure 1.**
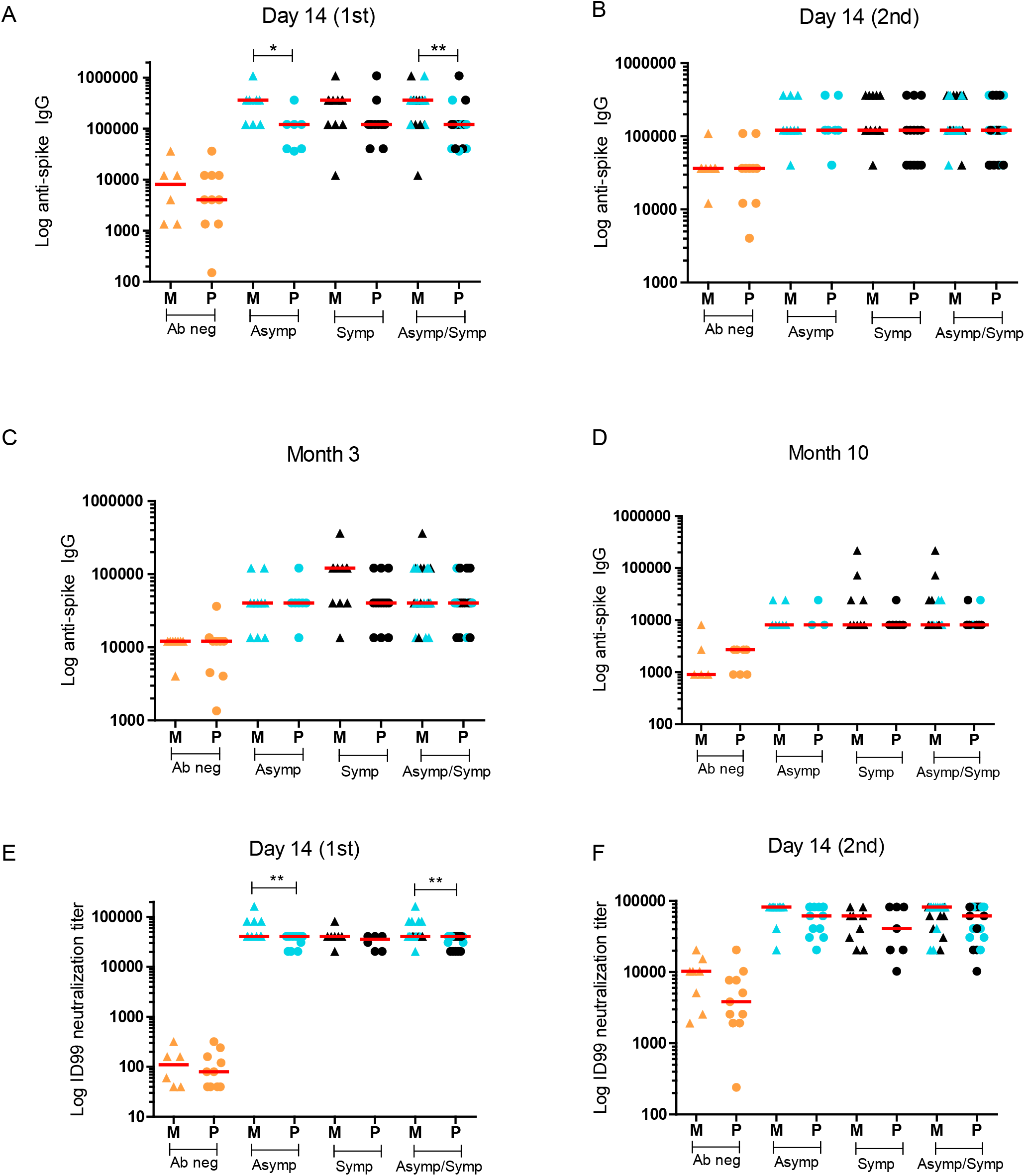
Comparison of plasma binding and neutralization titers by vaccination group. Antibody negative, Asymptomatic, Symptomatic, and SARS2-infected (combined Asymptomatic/Symptomatic) groups shown in orange, blue and black colors, respectively. Y-axis represents reciprocal endpoint IgG titer (or ID99 neutralization titer) to SARS-CoV-2 spike trimer in a logarithmic scale. Intra-group binding and neutralization titers at the different time points were analyzed by two-tailed Mann-Whitney. M= Moderna (triangles); P= Pfizer (circles). * = P ≤ 0.05; ** = P ≤ 0.01; *** = P ≤ 0.001; **** = P ≤ 0.0001

**Supplementary Figure 2.**
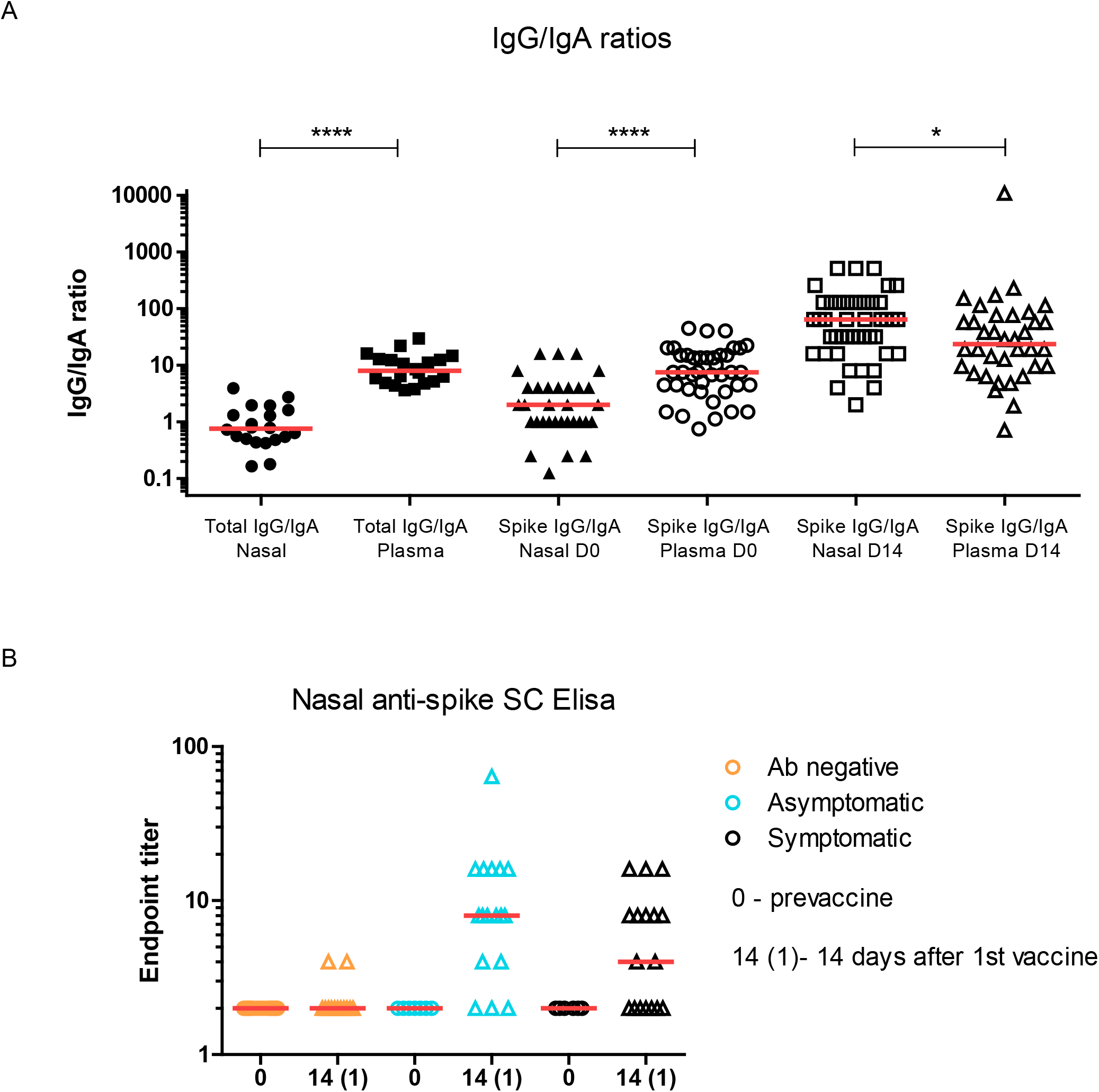
Nasal IgG/IgA ratio, and plasma IgG/IgA ratios, and nasal secretory antibody. A) IgG/IgA ratios measured in nasal samples and plasma samples in combined Asymptomatic and Symptomatic groups. Total IgG/IgA measured in nasal and plasma samples from various time points, and total IgG/IgA and anti-spike trimer IgG/IgA measured at baseline and Day 14 after the first vaccination. B) Anti-spike antibody carrying the secretory protein was measured in samples at baseline and 14 days after the first vaccination. For calculations IgG/IgA, sample pairs where both values were below limit of detection were excluded. Horizontal red lines represent median values. * = P ≤ 0.05; ** = P ≤ 0.01; *** = P ≤ 0.001; **** = P ≤ 0.0001

**Supplemental Figure 3.**
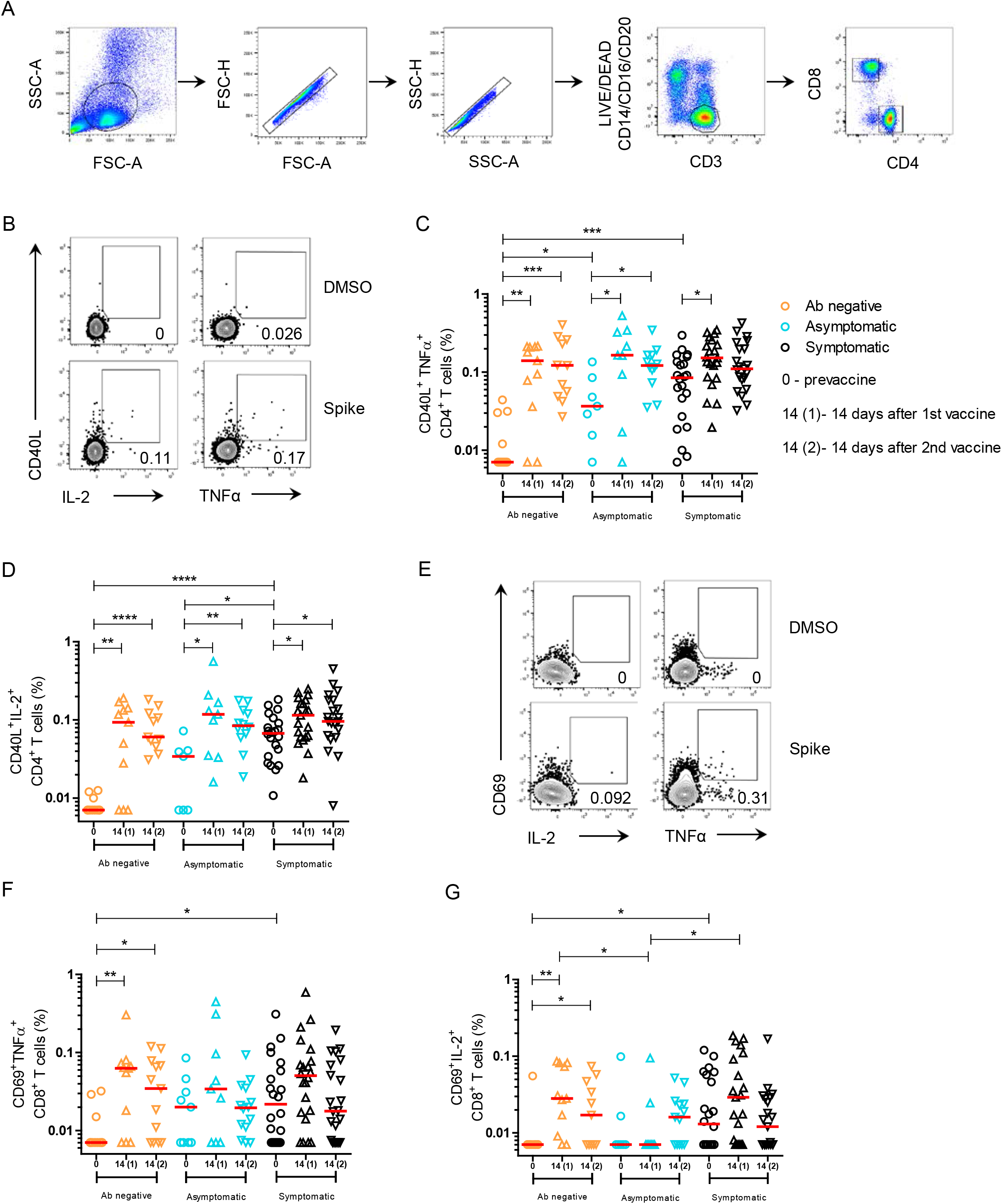
Spike-specific CD4^+^ and CD8^+^ T cells. **(A)** Gating strategy to define CD4^+^ and CD8^+^ T cells for AIM and ICS assays. **(B)** Representative gating strategy of CD40L^+^ IL-2^+^ and CD40L^+^ TNFα^+^ CD4^+^ T cells stimulated with spike peptide pool or unstimulated (DMSO). **(C)** Spike-specific CD40L^+^ TNFα ^+^ CD4^+^ T cells of total CD4^+^ T cells. **(D)** Spike-specific CD40L^+^ IL-2 ^+^ CD4^+^ T cells of total CD4^+^ T cells. **(E)** Representative gating strategy of CD69^+^ IL-2^+^ and CD69^+^ TNFα^+^ CD8^+^ T cells. **(F)** Spike-specific CD69^+^ TNFα ^+^ CD8^+^ T cells of total CD8^+^ T cells. **(G)** Spike-specific CD69^+^ IL-2^+^ CD8^+^ T cells of total CD8^+^ T cells. All data are background subtracted. Horizontal red lines represent median values.

**Supplemental Figure 4.**
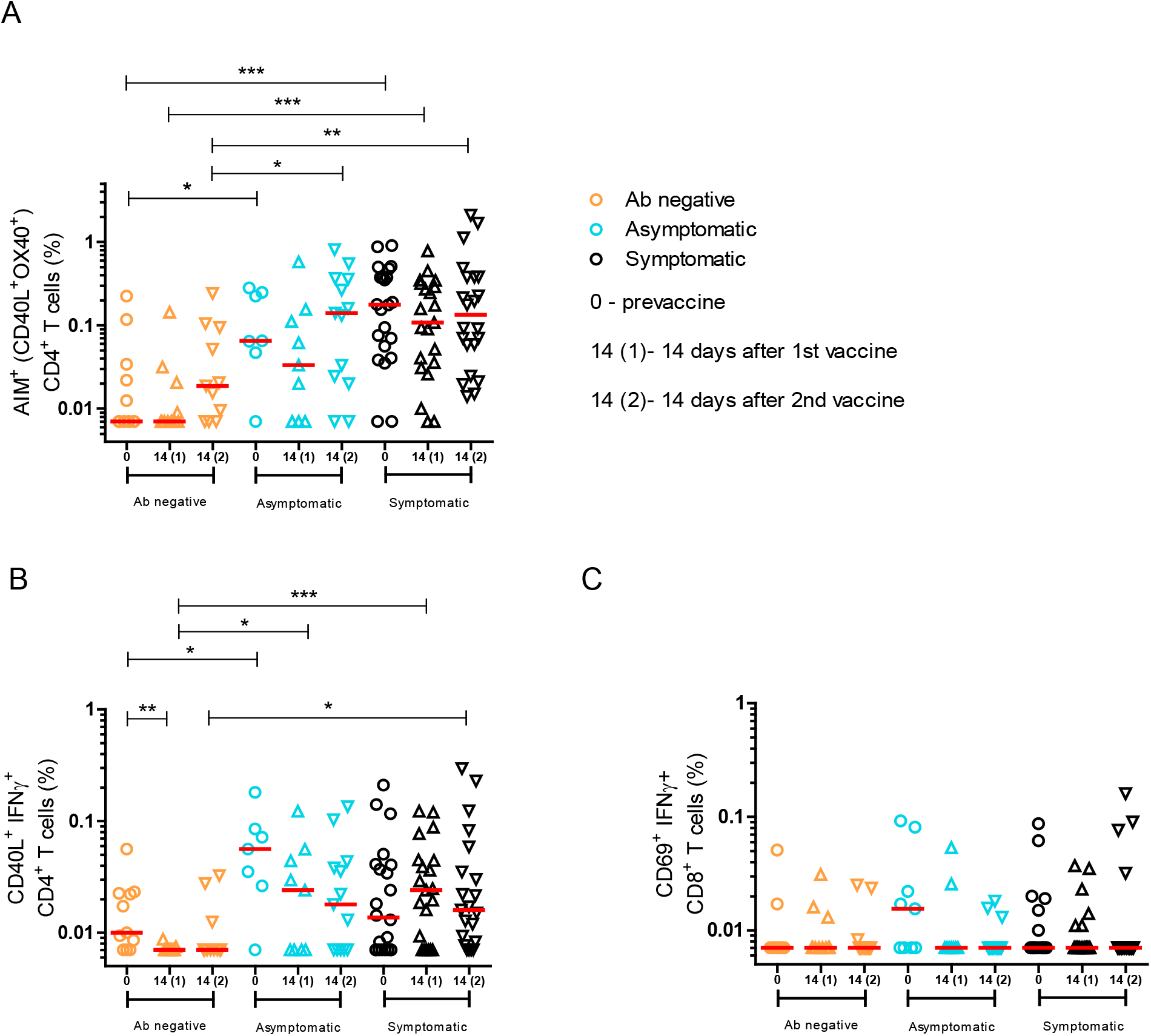
Non-spike-specific CD4^+^ and CD8^+^ T cells. **(A)** Non-spike-specific AIM^+^ (surface CD40L^+^ OX40^+^) CD4^+^ T cells of total CD4^+^ T cells. **(B)** Non-spike-specific CD40L^+^ IFNγ ^+^ CD4^+^ T cells of total CD4^+^ T cells. **(C)** Non-spike-specific CD69^+^ IFNγ ^+^ CD8^+^ T cells of total CD8^+^ T cells following stimulation with MP_R. All data are background subtracted. Horizontal red lines represent median values.

**Supplemental Figure 5.**
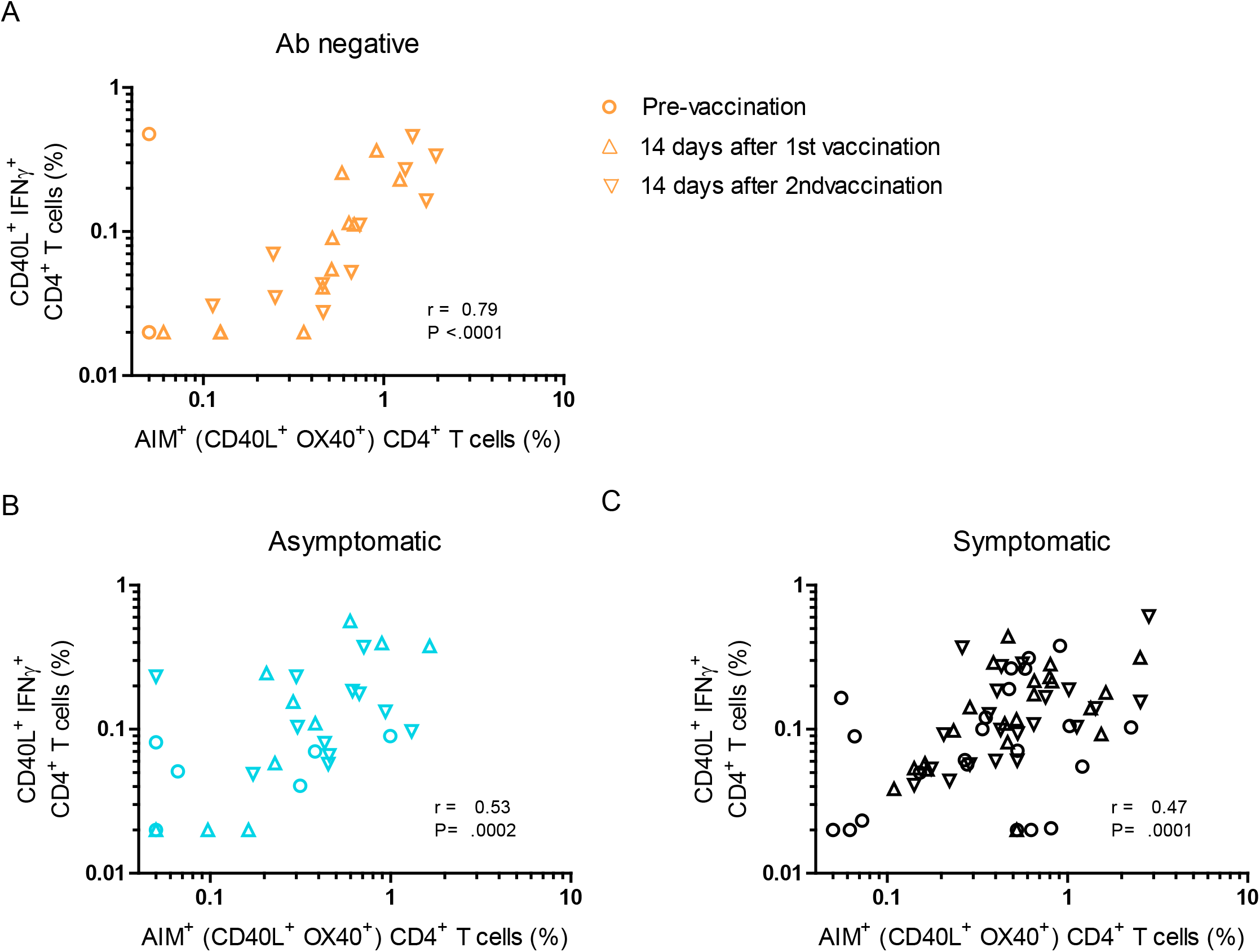
Correlation between spike-specific AIM^+^ and ICS^+^ CD4^+^ T cells. Correlation between AIM^+^ (surface CD40L^+^ OX40^+^) and CD40L^+^ IFNγ^+^ CD4^+^ T cells in **(A)** Antibody negative and **(B, C)** SARS2 exposed individuals. Statistics were calculated using Spearman’s correlation.

**Supplementary Figure 6.**
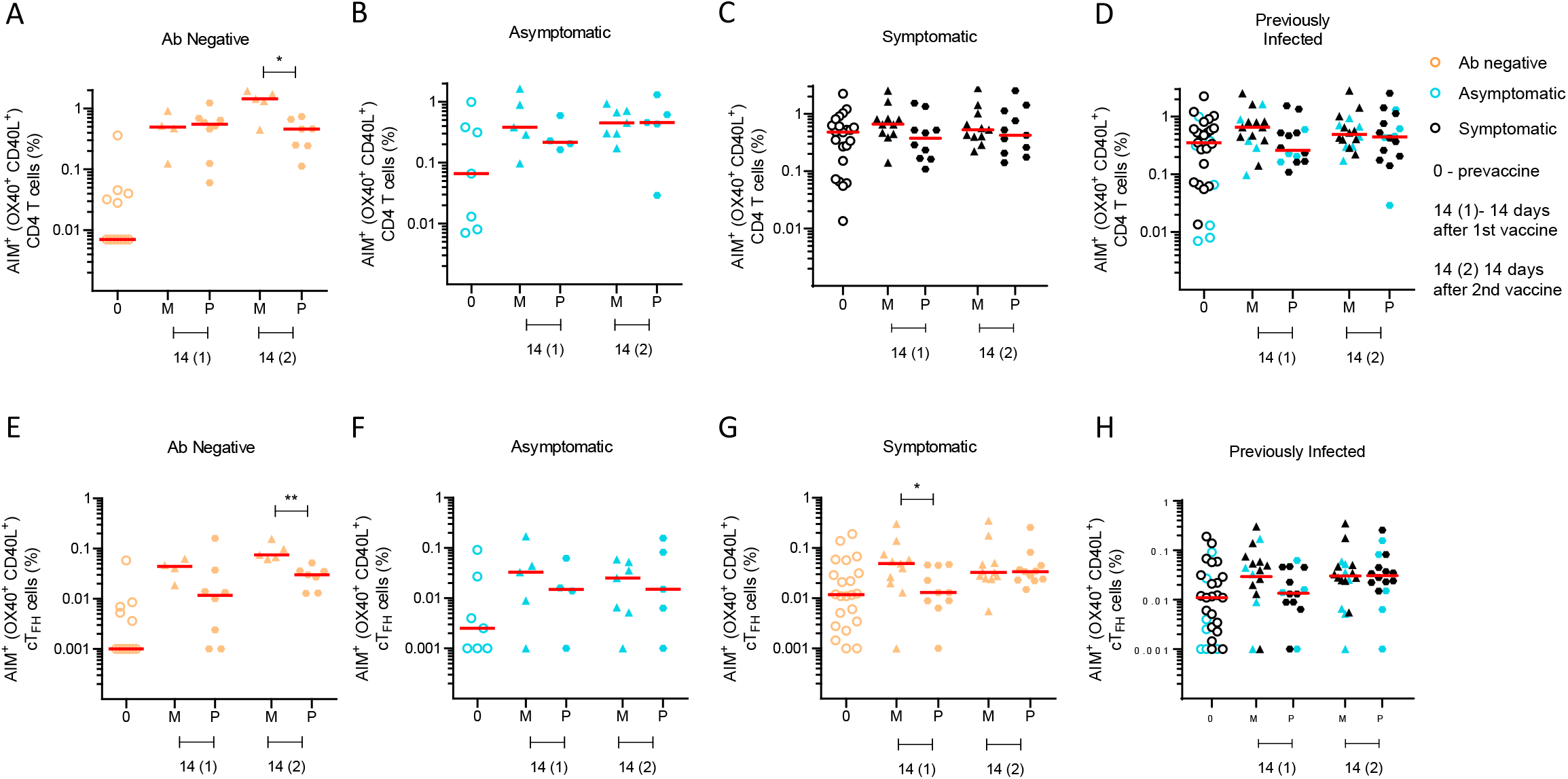
Spike-specific CD4^+^ and cT_FH_ cells detected by AIM assay in Moderna and Pfizer recipients. AIM^+^ (surface CD40L^+^ OX40^+^) spike-specific CD4^+^ T cells and cT_FH_ cells of total CD4^+^ T cells in (A,E) Antibody negative (orange), (B,F) Asymptomatic (blue), (C,G) Symptomatic (black) and (D,H) SARS2-infected (combined Asymptomatic/Symptomatic) groups. M= Moderna; P= Pfizer. Horizontal red lines represent median.

**Supplementary Figure 7.**
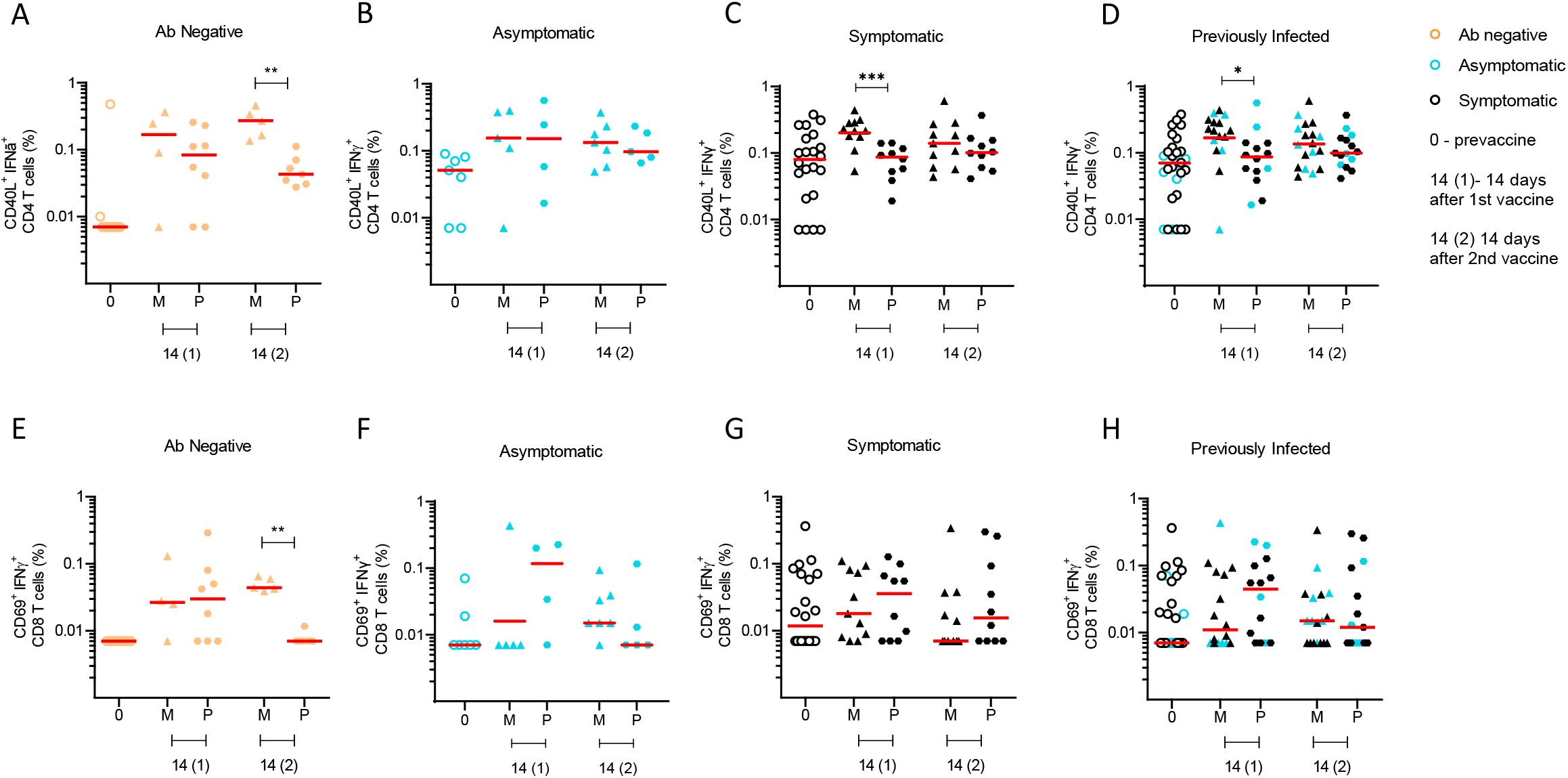
Spike-specific CD4^+^ and CD8^+^ cells detected by ICS assay in Moderna and Pfizer recipients. Spike-specific CD40L^+^ IFNγ^+^ CD4^+^ T cells of total CD4^+^ T cells in (A,E) Antibody negative (orange), (B,F) Asymptomatic (blue), (C,G) Symptomatic (black) and (D,H) SARS2-infected (combined Asymptomatic/Symptomatic) groups. Spike-specific CD69^+^ IFNγ^+^ CD8^+^ T cells of total CD8^+^ T cells in (E) Antibody negative, (F) Asymptomatic, (G) Symptomatic and (H) Previously infected (Asymptomatic/Symptomatic) groups. M= Moderna; P= Pfizer. Horizontal red lines represent median.

**Supplemental Figure 8.**
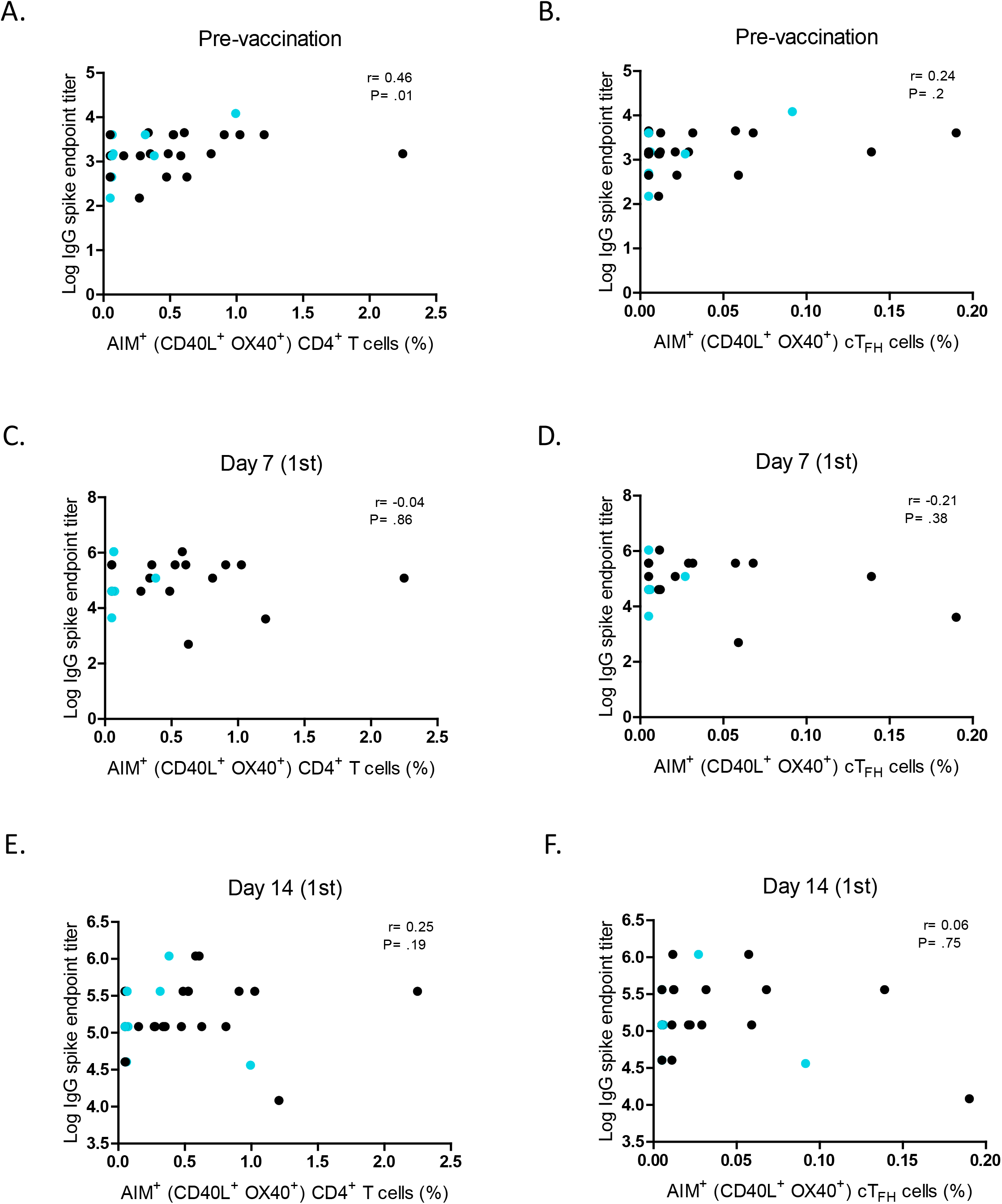
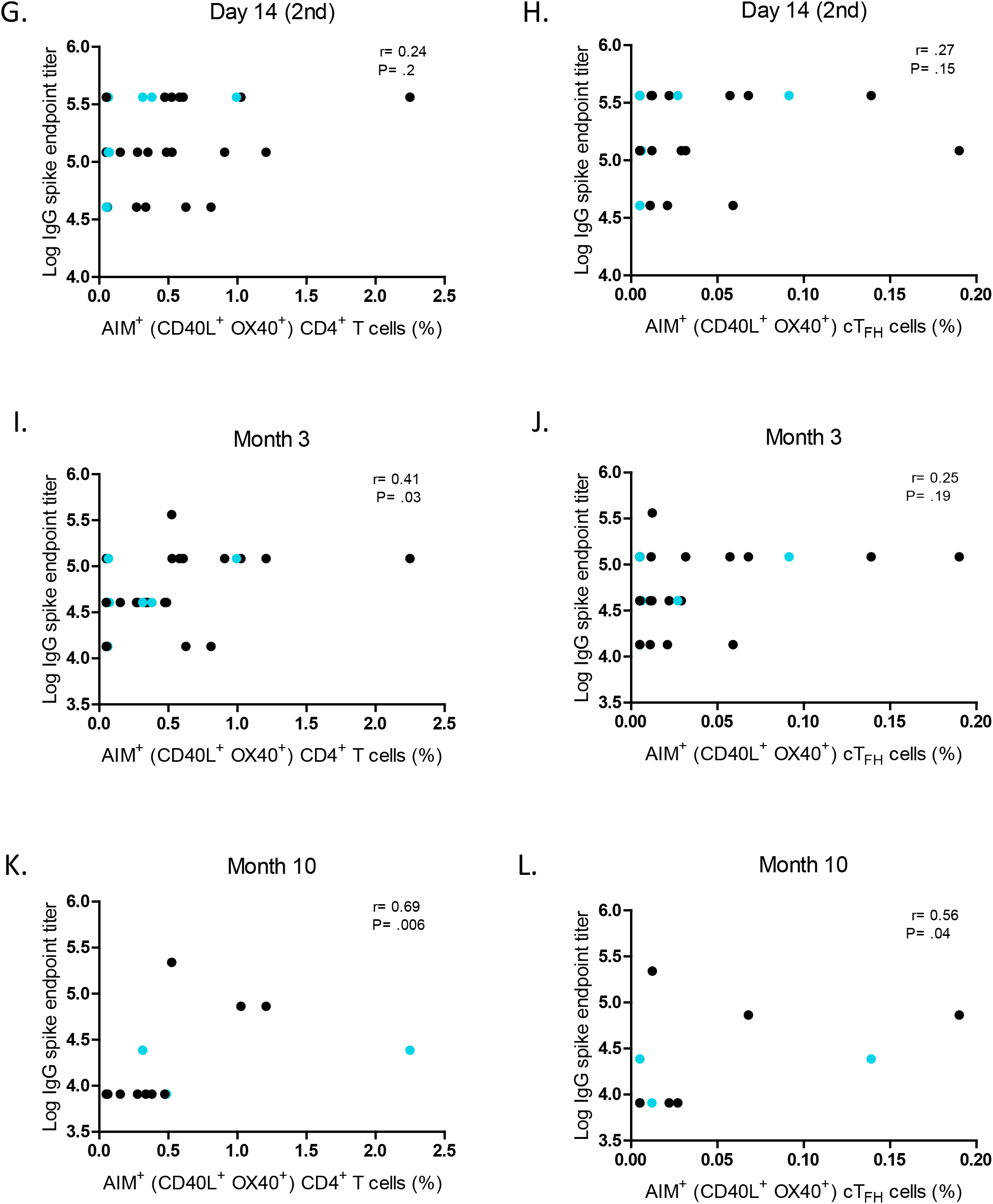
Correlation between pre-existing spike-specific CD4^+^ T cells and Spike IgG. Correlation between pre-existing spike-specific AIM^+^ (surface CD40L^+^ OX40^+^) CD4^+^ T cells (panels A,C,E,G,I,K) or AIM^+^ (surface CD40L^+^ OX40^+^) cT_FH_ cells (panels B,D,F,H,J,K) and spike IgG prior to vaccination, 7 days after 1^st^ vaccination,14 days after 1^st^ vaccination, 14 days after 2^nd^ vaccination, 3 months after 2^nd^ vaccination and 10 months after 2^nd^ vaccination. Asymptomatic, and Symptomatic groups shown in blue and black circles, respectively. Statistics were calculated using Spearman’s correlation.

**Supplemental Figure 9.**
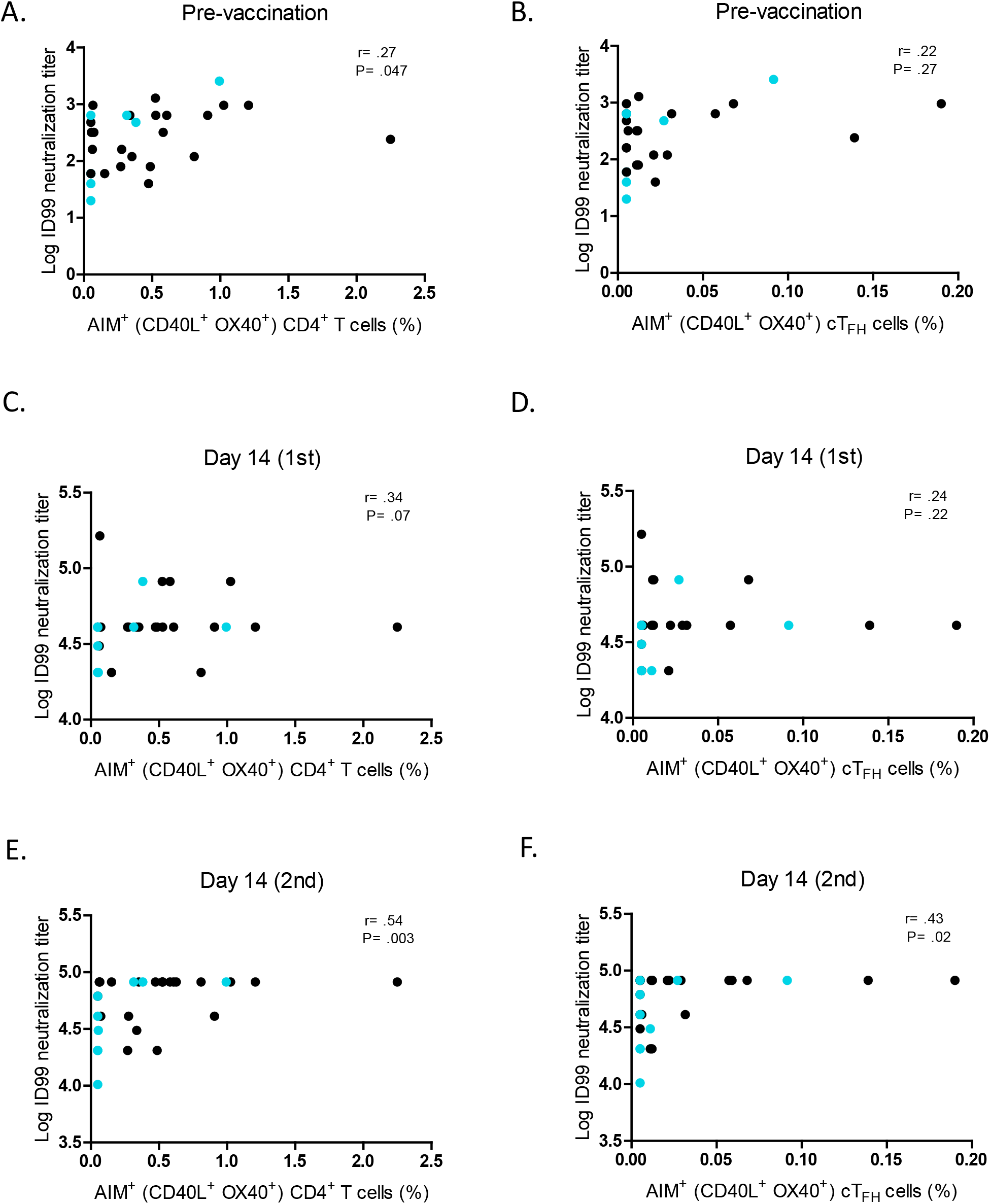
Correlation between pre-existing spike-specific CD4^+^ T cells and neutralization. Correlation between pre-existing spike-specific AIM^+^ (surface CD40L^+^ OX40^+^) CD4^+^ T cells (panels A,C,E) or AIM^+^ (surface CD40L^+^ OX40^+^) cT_FH_ cells (panels B,D,F) and ID99 neutralization prior to vaccination, 14 days after 1^st^ vaccination, and 14 days after 2^nd^ vaccination. Asymptomatic and Symptomatic groups shown in blue and black circles, respectively. Statistics were calculated using Spearman’s correlation.

## References

1. A. M. Cavanaugh, K. B. Spicer, D. Thoroughman, C. Glick, K. Winter, Reduced Risk of Reinfection with SARS-CoV-2 After COVID-19 Vaccination - Kentucky, May-June 2021. MMWR Morb Mortal Wkly Rep 70, 1081–1083 (2021).

2. K. B. Pouwels et al., Effect of Delta variant on viral burden and vaccine effectiveness against new SARS-CoV-2 infections in the UK. Nat Med, (2021).

3. S. J. Thomas et al., Safety and Efficacy of the BNT162b2 mRNA Covid-19 Vaccine through 6 Months. N Engl J Med 385, 1761–1773 (2021).

4. K. E. Mullins et al., Validation of COVID-19 serologic tests and large scale screening of asymptomatic healthcare workers. Clin Biochem, (2021).

5. E. G. Levin et al., Waning Immune Humoral Response to BNT162b2 Covid-19 Vaccine over 6 Months. N Engl J Med, (2021).

6. Y. Goldberg et al., Waning Immunity after the BNT162b2 Vaccine in Israel. N Engl J Med, (2021).

7. S. Nanduri et al., Effectiveness of Pfizer-BioNTech and Moderna Vaccines in Preventing SARS-CoV-2 Infection Among Nursing Home Residents Before and During Widespread Circulation of the SARS-CoV-2 B.1.617.2 (Delta) Variant - National Healthcare Safety Network, March 1-August 1, 2021. MMWR Morb Mortal Wkly Rep 70, 1163–1166 (2021).

8. S. Y. Tartof et al., Effectiveness of mRNA BNT162b2 COVID-19 vaccine up to 6 months in a large integrated health system in the USA: a retrospective cohort study. Lancet 398, 1407–1416 (2021).

9. F. Krammer et al., Antibody Responses in Seropositive Persons after a Single Dose of SARS-CoV-2 mRNA Vaccine. N Engl J Med 384, 1372–1374 (2021).

10. J. E. Ebinger et al., Antibody responses to the BNT162b2 mRNA vaccine in individuals previously infected with SARS-CoV-2. Nat Med 27, 981–984 (2021).

11. K. Abu Jabal et al., Impact of age, ethnicity, sex and prior infection status on immunogenicity following a single dose of the BNT162b2 mRNA COVID-19 vaccine: real-world evidence from healthcare workers, Israel, December 2020 to January 2021. Euro Surveill 26, (2021).

12. R. Levi et al., One dose of SARS-CoV-2 vaccine exponentially increases antibodies in individuals who have recovered from symptomatic COVID-19. J Clin Invest 131, (2021).

13. S. Saadat et al., Binding and Neutralization Antibody Titers After a Single Vaccine Dose in Health Care Workers Previously Infected With SARS-CoV-2. JAMA 325, 1467–1469 (2021).

14. E. Fraley et al., Humoral immune responses during SARS-CoV-2 mRNA vaccine administration in seropositive and seronegative individuals. BMC Med 19, 169 (2021).

15. L. Stamatatos et al., mRNA vaccination boosts cross-variant neutralizing antibodies elicited by SARS-CoV-2 infection. Science, (2021).

16. E. Andreano et al., Hybrid immunity improves B cells and antibodies against SARS-CoV-2 variants. Nature, (2021).

17. Z. Wang et al., Naturally enhanced neutralizing breadth against SARS-CoV-2 one year after infection. Nature 595, 426–431 (2021).

18. R. R. Goel et al., Distinct antibody and memory B cell responses in SARS-CoV-2 naive and recovered individuals following mRNA vaccination. Sci Immunol 6, (2021).

19. C. Manisty et al., Antibody response to first BNT162b2 dose in previously SARS-CoV-2-infected individuals. Lancet 397, 1057–1058 (2021).

20. K. A. Callow, Effect of specific humoral immunity and some non-specific factors on resistance of volunteers to respiratory coronavirus infection. J Hyg (Lond) 95, 173–189 (1985).

21. G. M. Spiekermann et al., Receptor-mediated immunoglobulin G transport across mucosal barriers in adult life: functional expression of FcRn in the mammalian lung. J Exp Med 196, 303–310 (2002).

22. D. S. Khoury et al., Neutralizing antibody levels are highly predictive of immune protection from symptomatic SARS-CoV-2 infection. Nat Med 27, 1205–1211 (2021).

23. S. Feng et al., Correlates of protection against symptomatic and asymptomatic SARS-CoV-2 infection. Nat Med 27, 2032–2040 (2021).

24. Z. Rikhtegaran Tehrani et al., Performance of nucleocapsid and spike-based SARS-CoV-2 serologic assays. PLoS One 15, e0237828 (2020).

25. C. Keech et al., Phase 1-2 Trial of a SARS-CoV-2 Recombinant Spike Protein Nanoparticle Vaccine. N Engl J Med 383, 2320–2332 (2020).

26. A. Grifoni et al., Targets of T Cell Responses to SARS-CoV-2 Coronavirus in Humans with COVID-19 Disease and Unexposed Individuals. Cell 181, 1489–1501 e1415 (2020).

27. J. Mateus et al., Low-dose mRNA-1273 COVID-19 vaccine generates durable memory enhanced by cross-reactive T cells. Science 374, eabj9853 (2021).

